# miR-28 plus ibrutinib as a novel combination therapy for Diffuse Large B Cell Lymphoma

**DOI:** 10.1101/2022.09.27.509662

**Authors:** Teresa Fuertes, Emigdio Álvarez-Corrales, Patricia Ubieto-Capella, Álvaro Serrano-Navarro, Carmen Gómez-Escolar, Antonio de Molina, Juan Méndez, Almudena R. Ramiro, Virginia G. de Yébenes

**Author notes:** Equal contribution (Almudena R. Ramiro, Virginia G. de Yébenes). **Corresponding authors**: Almudena R Ramiro. Centro Nacional de Investigaciones Cardiovasculares. Melchor Fernandez Almagro 3. 28029 Madrid. or Virginia G de Yebenes Universidad Complutense de Madrid. Pza. Ramón y Cajal, s/n. 28040 Madrid.

## Abstract

Diffuse large B cell lymphoma (DLBCL) is the most common aggressive B cell lymphoma and accounts for nearly 40% of cases of B cell non-Hodgkin lymphoma. DLBCL is generally treated with R-CHOP chemotherapy, but many patients do not respond or relapse after treatment. Here, we analyzed the therapeutic potential of the tumor suppressor microRNA-28 (miR-28) for DLBCL, alone and in combination with the Bruton’s tyrosine kinase inhibitor ibrutinib. Combination therapy with miR-28 plus ibrutinib potentiated the anti-tumor effects of monotherapy with either agent by inducing a specific transcriptional cell-cycle arrest program that impairs DNA replication. Moreover, we found that downregulation of the miR-28-plus-ibrutinib gene signature correlates with better survival of ABC-DLBCL patients. These results provide evidence for the effectiveness of a new miRNA-based ibrutinib combination therapy for DLBCL and unveil the miR-28-plus-ibrutinib gene signature as a new predictor of outcome in ABC-DLBCL patients.

**STATEMENT OF SIGNIFICANCE:** This study demonstrates that a miRNA-based combination therapy with ibrutinib is superior to both monotherapies in DLBCL. miR-28 plus ibrutinib combined therapy inhibits DLBCL growth through the induction of a transcriptional program that impairs DNA replication. Our results provide a new gene signature with prognostic value for ABC-DLBCL.

## INTRODUCTION

Diffuse large B cell lymphoma (DLBCL) is the most common aggressive lymphoma and accounts for nearly 40% of B cell non-Hodgkin lymphomas (B-NHL) [1]. DLBCL shows heterogeneous clinical manifestations and responses to therapy [1]. The disease was initially classified according to gene expression profiles into two major subtypes, called germinal center B cell-like (GCB) DLBCL and activated B cell-like (ABC) DLBCL [2]. Gene expression in GCB-DLCBL is characteristic of normal germinal center B cells, whereas ABC-DLBCL has post-germinal center features and expresses genes that are induced following B cell receptor (BCR) engagement and B cell activation. A major hallmark of ABC-DLBCL is activation of the NF-*k*B pathway [3]. Genetic profiling has provided a deeper understanding of the heterogeneous genetic aberrations in DLBCL, resulting in the identification of four DLBCL subtypes termed MCD, BN2, N1, and EZB, which differ not only in their genetic alterations but also in their gene expression signatures and clinical outcomes [4].

About 60% of DLBCL patients are cured by chemotherapy with rituximab plus cyclophosphamide, doxorubicin, vincristine, and prednisone (R-CHOP) [5] [6]. However, ABC-DLBCL has a worse prognosis than GCB-DLBCL [2, 7], and disease progression after R-CHOP chemotherapy is prevented in only 40% of ABC-DLBCL patients [1, 7]. Moreover, most patients who relapse or are refractory to initial therapy succumb to their disease [8]. New drug development for DLBCL patients over the past decade has focused on identifying specific, more effective, and non-cytotoxic targeted therapies [9] [1]. A major advance in DLBCL therapy development can be achieved with the advent of ibrutinib, a Bruton’s tyrosine kinase (BTK) inhibitor that blocks B-cell receptor (BCR) dependent NF-*k*B activation and is approved for the treatment of mantle cell lymphoma, chronic lymphocytic leukemia, marginal zone lymphoma and Waldenström’s macroglobulinemia [10] [11]. Clinical trials have shown that ibrutinib is especially effective when used in combination with R-CHOP to treat young ABC-DLBCL patients of the MCD and N1 genetic subtypes [12] [13]. However, DLBCL patients are often elderly and have age-related comorbidities, and therefore there is a need for new non-chemotherapy-based combination regimens that enhance the response to ibrutinib.

microRNAs (miRNAs) are non-coding RNAs that negatively regulate gene expression. Through imperfect base pairing, miRNAs bind to their target mRNAs and promote their degradation or translational blockade. Each miRNA binds numerous target mRNAs, regulating the expression of gene networks [14]. miRNA expression is frequently dysregulated in cancer, and miRNAs have been shown to play a causative role in B cell lymphoma development, acting as tumor suppressors or tumor promoting OncomiRs [15]. miRNA-based therapeutics is envisioned as an attractive alternative approach for B-NHL and other cancers (reviewed in [16] and [17]). We previously reported that miR-28 is a tumor suppressor miRNA that regulates a BCR-dependent signaling network in B cells [18, 19]. Here, we have assessed the therapeutic value of miR-28 for DLBCL alone and in combination with ibrutinib. We present evidence that miR-28 enhances the anti-tumor effect of ibrutinib, promoting a specific gene expression program that inhibits DNA replication and cell proliferation in DLBCL.

## RESULTS

### miR-28 inhibits DLBCL tumor growth

To evaluate the anti-tumor activity of miR-28 in treatment-refractory B cell lymphoma, we used different DLBCL models. Using human DLBCL xenografts, we first assessed the anti-tumor activity of a synthetic miR-28 analog (mimic) [20]. Three GCB-DLBCL (DOHH2, OCI LY19, and DB) and two ABC-DLBCL (MD-901 and Riva) human cell lines were injected subcutaneously into NOD *scid* gamma (NSG) mice. Tumors were allowed to reach at least 150 mm^3^, and mice were then treated with intratumoral injections of 0.5 nmol miR-28 or control mimics (Figure S1A). The synthetic miR-28 mimic significantly inhibited tumor growth of all five human DLCBL cell lines tested, with similar efficacy in the ABC-DLBCL and GCB-DLBCL cell lines (Figure 1A, 1B). We next used a patient-derived xenograft (PDX) ABC-DLBCL model from the PRoXe xenograft repository [21]. This PDX was isolated from an untreated patient and is characterized by the expression of human CD19, CD20, and IgM and by high expression of the Ki-67 proliferation marker when injected into NSG mice (Figure 1C). The synthetic miR-28 mimic significantly inhibited the tumor growth of DLCBL PDX xenografts after two administrations (Figure 1D, S1B). RNA-Seq transcriptome analysis of ABC-DLCBL PDX tumors 5 days after treatment with the scramble miRNA (control) or the miR-28 mimic (3 tumors per treatment) revealed miR-28–induced expression changes in 314 protein-coding transcripts (Figure 1E, Table S1A). Ingenuity Pathway Analysis (IPA) showed that miR-28 coordinately inhibited the expression of gene networks implicated in tumor growth (Figure 1F,Table S1B). In addition, Gene Set Enrichment Analysis (GSEA) showed that miR-28 significantly downregulated transcripts known to be overexpressed in DLBCL patients (p=0.026, NES=-1.13) (Figure 1G). Together, these results show that synthetic miR-28 analogs inhibit DLBCL growth *in vivo* through the downregulation of gene networks involved in tumor growth.

**Figure 1.**
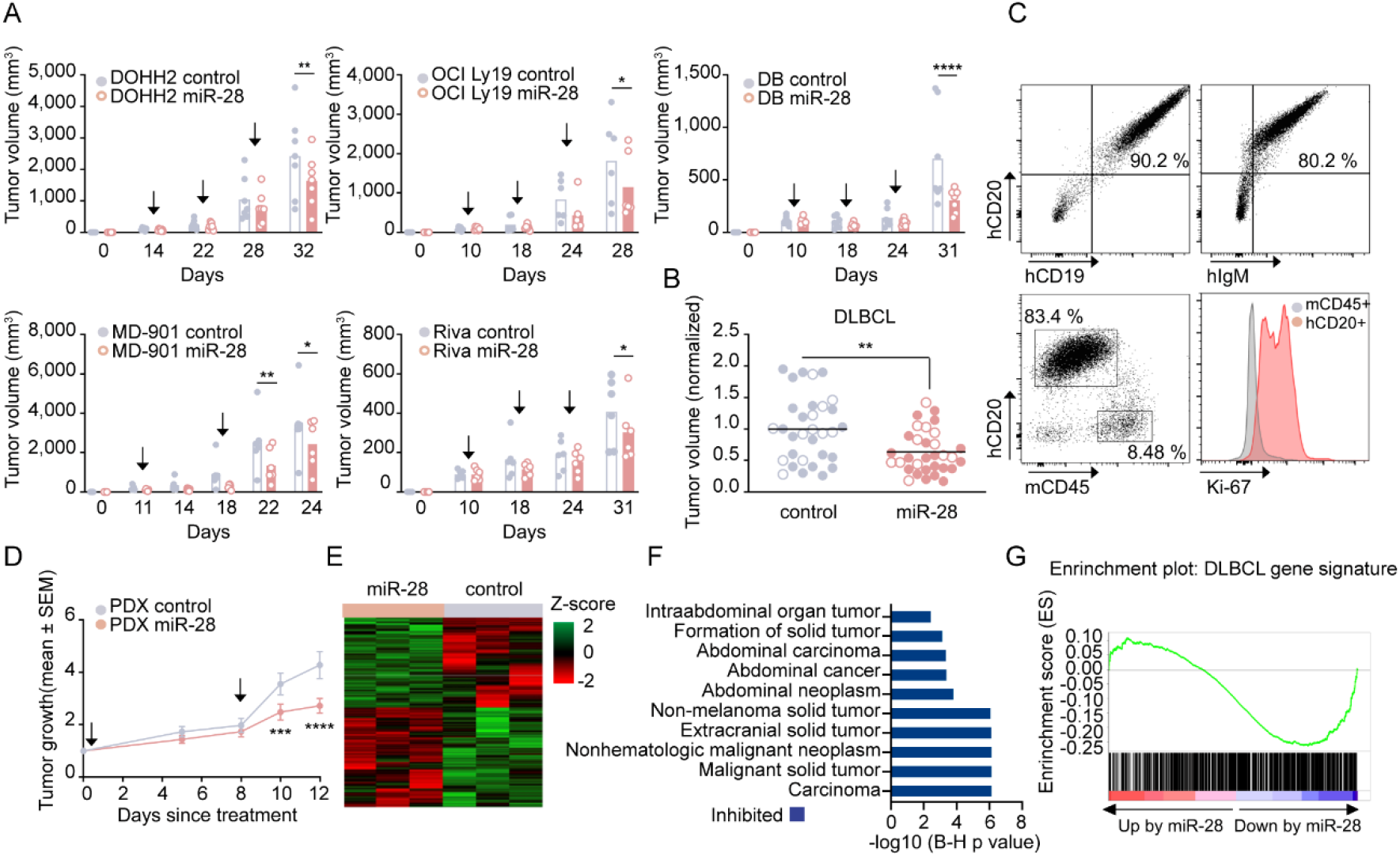
miR-28 inhibits DLBCL tumor growth. **(A-B)** Tumor growth of GCB-DLBCL and ABC-DLBCL cell-line xenografts in NSG mice after intratumoral treatment with 0.5 nmol miRNA mimics: miR-28 (pink) or scramble control (gray). **(A)** GCB-DLBCL xenografts (upper graphs): DOHH2 (n=7), OCIL Ly19 (n=6), DB (n=7). ABC-DLBCL xenografts (bottom graphs): MD-901 (n=6), Riva (n=6). Bar plots show tumor volume at the indicated time points. Arrows indicate dosing schedules. *P<0.05, **P<0.01, ****P<0.0001, linear mixed model. **(B)** End-point tumor volume normalized to the mean volume in each control group for the xenografts shown in A (ABC-DLBCL, open circles; GCB-DLBCL, filled circles). **P<0.01, unpaired *t* test. **(C)** Representative flow cytometry plots of spleen-derived PDX cells (CD20+, CD19+, IgM+, mouse CD45, Ki-67+) 26 to 31 days after infusion of PDX cells in NSG mice. **(D-F)** Effect of miR-28 on tumor growth in a DLBCL–patient-derived xenograft (PDX) mouse model (PRoXe DFBL-18689-V2). **(D)** ABC-DLBCL PDX tumors were treated with intratumoral injections of 0.5 nmol miR-28 mimics or control (n=24 tumors, 3 independent experiments). Tumor growth was calculated as tumor volume at the indicated time points normalized to the initial tumor volume. ***P<0.001, ****P<0.0001, linear mixed model. **(E)** RNA-Seq analysis of ABC-DLBCL PDX tumors 5 days after a single miRNA mimic treatment (n=3 miR-28, n=3 control). The heatmap shows expression of the 314 genes found to be differentially expressed (ANOVA p-value <0.05). **(F)** The graph shows the top 10 significantly changed Pathways identified by Ingenuity Pathway Analysis (IPA). Pathway activity prediction is shown with a color code (blue, inhibited by miR-28). P-values were corrected for multiple testing using the Benjamini-Hochberg (B–H) false discovery rate. **(G)** Gene Set Enrichment Analysis (GSEA) of a DLBCL gene signature [42]in miR-28-treated PDX tumors (FWER p-value = 0.025, FDR q value= 0.066, NES= −1.13).

### miR-28 potentiates the anti-tumor effect of ibrutinib

We next investigated whether miR-28 can sensitize DLBCL to the BTK inhibitor ibrutinib. We first tested *in vitro* cell viability and growth in the presence of ibrutinib after the induction of miR-28 expression in four human DLBCL cell lines. Three ABC-DLBCL (MD-901, U2932 and HBL1) cell lines and one GCB-DLBCL (SUDHL4) cell line were transduced with a doxycycline-inducible lentiviral vector encoding miR-28 (pTRIPZ-miR-28) or a scramble sequence (pTRIPZ-scramble) together with red fluorescent protein (RFP) (Figure S2A). We first tested ibrutinib sensitivity in each control-transduced cell line, as analyzed 3 days after ibrutinib treatment. Cell viability was quantified as the total number of viable cells measured by CellTiter-Glo® luminescent ATP detection assay (Figure 2A) and as the proportion of live cells unstained with the DAPI DNA-dye, detected by flow cytometry (Figure S2B). As expected, the highest sensitivity was detected in HBL1 cells, which harbor MYD88 L265P and CD79B mutations that confer high sensitivity to ibrutinib [22]. In all four DLBCL cell lines tested, doxycycline-induced miR-28 expression reduced the number of viable cells upon ibrutinib treatment (Figure 2A, S2B). The enhancement of ibrutinib anti-tumor activity in the presence of miR-28 differed among the different cell lines, with the strongest effect detected in MD-901 cells (Figure 2A, S2B). We next assessed the ability of miR-28 to enhance the anti-tumor effect of ibrutinib on the growth of DLBCL xenografts *in vivo*. MD-901 cells transduced with pTRIPZ-miR-28 or pTRIPZ-scramble were injected subcutaneously into NSG mice, and once tumors were detectable, mice were given doxycycline in the drinking water for 9 days to induce miR-28 or scramble expression. During the same period, mice received daily intraperitoneal injections of ibrutinib (40 mg/kg) or vehicle (DMSO). There were thus four treatment groups: scramble alone (control); scramble plus ibrutinib; miR-28 alone; and miR-28 plus ibrutinib combination therapy. Induced miR-28 expression significantly enhanced the inhibitory effect of ibrutinib on MD-901 ABC-DLBCL tumor volume, growth, and weight (Figure 2B, Table S2). Together, these results demonstrate that miR-28 enchances DLBCL sensitivity to ibrutinib.

**Figure 2.**
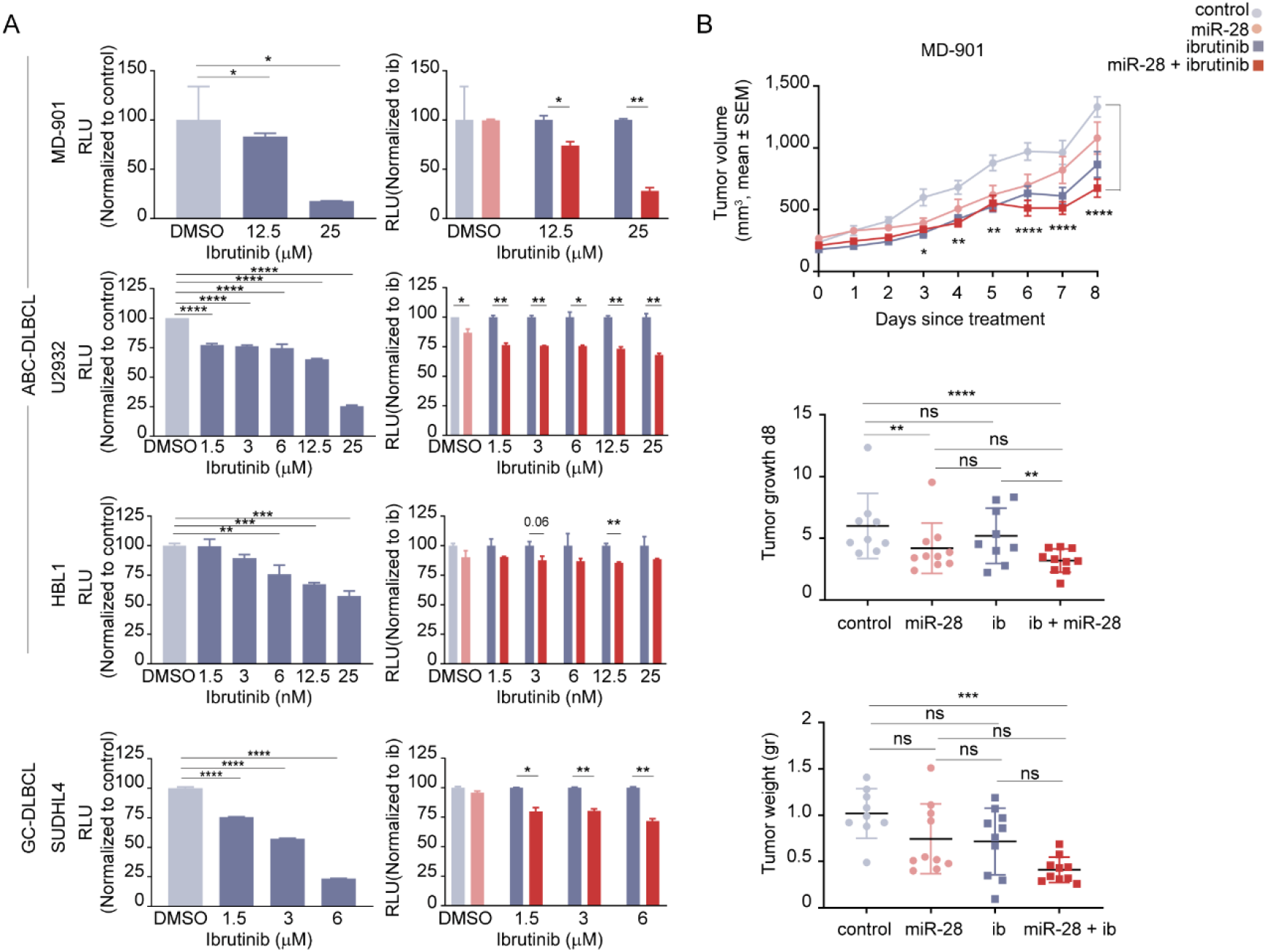
miR-28 potentiates the anti-tumor effect of ibrutinib against DLBCL. **(A)** Effect of miR-28 on the growth of DLBCL cell lines treated with ibrutinib *in vitro*. DLBCL cell lines transduced with pTRIPZ-miR-28 (miR-28) or pTRIPZ-scramble (control) were treated with the indicated concentrations of ibrutinib in the presence of doxycycline. Viable cell number was determined at day 3 with CellTiter-Glo® reagent. Each row represents a different cell line: ABC-DLBCL subtype (upper rows: MD-901, U2932 and HBL1); GCB-DLBCL subtype (lower row SUDHL4). Error bars denote SD of duplicates. Left bar charts show sensitivity to ibrutinib in control cells, calculated as Relative Luminescence Units (RLU) normalized to control (control DMSO-treated, light gray; ibrutinib-treated, dark gray). Right bar charts show the effect of treatment with miR-28 plus ibrutinib (dark red) normalized to control ibrutinib treated cells (dark gray). miR-28 cells normalized to control are shown in light pink. *P<0.05, **P<0.01, ***P<0.001, ****P<0.0001, unpaired *t* test. **(B)** Effect of miR-28+ibrutinib combination therapy on ABC-DLBCL xenografts growth *in vivo*. Xenograft growth from MD-901 cells transduced with doxycycline-inducible pTRIPZ-miR-28 or pTRIPZ-scramble in NSG mice treated with daily intraperitoneal injections of ibrutinib or vehicle and administered with doxycycline in the drinking water: control (light gray circles), miR-28 (light pink circles), ibrutinib (dark gray squares), ibrutinib+miR-28 (dark red squares). The top graph shows tumor volume (mean volume ± SEM; n=5 mice and 10 tumors per group). *P<0.05, **P<0.01, ****P<0.0001 vs control, linear mixed model). The middle graph shows tumor growth, calculated as tumor volume at day 8 normalized to initial tumor volume **P<0.01, ****P<0.0001, linear mixed model. See Table S2 for a complete statistical analysis. The bottom graph shows tumor weight at experimental endpoint (day 9 after treatment initiation). ***P<0.001, one-way ANOVA.

### miR-28+ibrutinib combination therapy induces a specific transcriptional program that inhibits the cell cycle and DNA replication

To investigate the effect of miR-28+ibrutinib combination therapy on the DLBCL transcriptome, we conducted an RNA-Seq analysis to compare gene expression profiles after monotherapy versus combined miR-28+ibrutinib treatment. MD-901 cells transduced with pTRIPZ-miR-28 or pTRIPZ-scramble were induced with doxycycline for 24 hours before ibrutinib treatment. RNA-Seq was performed on RFP+ cells isolated by flow cytometry after 0, 4, and 20 hours of ibrutinib exposure (Figure S3A). Gene expression changes were identified in response to miR-28 expression, ibrutinib treatment, and miR-28+ibrutinib combination (Figure S3C, Table S3). This analysis identified a set of 157 genes (adjusted p-value <0.05) that were significantly altered exclusively in MD-901 cells treated with miR-28+ibrutinib for 20h, which we termed “miR-28+ib signature” (Figure 3A, S3B and Table S4A). Genes in the miR-28+ib signature were mostly associated with DNA replication, DNA repair, and cell cycle pathways, and miR-28+ibrutinib combination therapy mostly resulted in their downregulation (Figure 3A). Consistent with these findings, Ingenuity Pathway analysis and GSEA showed that miR-28+ibrutinib combination therapy coordinately inhibited the activity of gene networks involved in the cell cycle and chromosomal replication (Figure 3B, S3D and Table S4B-C). In addition, STRING functional enrichment analysis identified a protein-protein interaction network composed of 65 proteins related to cell division whose expression was predominantly downregulated upon miR-28+ibrutinib combination therapy (Figure 3C).

**Figure 3.**
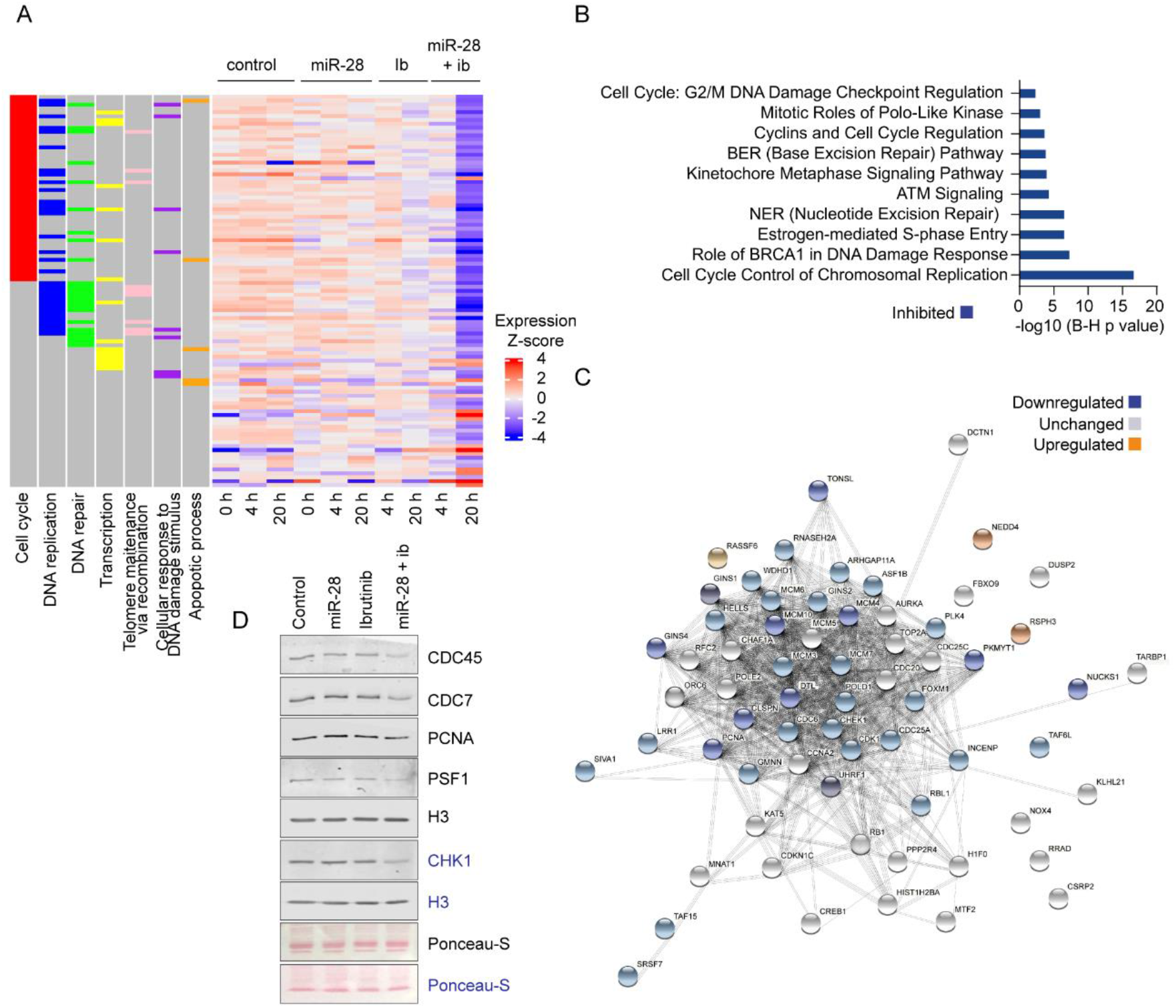
Effect of miR-28 plus ibrutinib combined treatment on the DLBCL-cell transcriptome. Doxycycline-induced MD-901 cells transduced with pTRIPZ-miR-28 or pTRIPZ-control were treated with 25 μM ibrutinib. Whole transcriptome RNA-Seq was performed in RFP^+^ cells isolated by flow cytometry after 0, 4, and 20 hours of ibrutinib treatment. **(A)** Heatmap showing Z-score expression values for the 157 miR-28+ib signature genes at 0, 4 and 20 hours in the four treatment conditions (control, miR-28, ibrutinib, and miR-28+ibrutinib). 114 genes were downregulated in the miR-28+ibrutinib condition. Columns on the left depict the 7 most representative Gene Ontology terms in the miR-28+ib signature. **(B)** Bar plot showing Benjamini-Hochberg (B-H) adjusted p-values (q val) for the top 10 Signaling Pathways in the miR-28+ib signature. Pathway activity prediction is shown with a color code (blue: inhibited by miR-28+ibrutinib combined treatment). **(C)** STRING protein-protein interaction network identified in the miR-28+ib signature by IPA. Upregulated proteins are depicted in orange, downregulated proteins in blue, and unchanged proteins in white. **(D)** Western blot analysis of selected replication proteins in doxycycline-induced pTRIPZ-miR-28− or pTRIPZ-scramble–transduced MD-901 cells after treatment for 20 hours with 25 μM ibrutinib or DMSO.

Within this DNA replication and cell division network, we analyzed the expression of a selected set of proteins required for the formation and activation of DNA replication origins: 1) proteins of the pre-initiation complex (pre-IC) required for helicase activation and replisome assembly during the G1-S transition, including CDC45, CDC7, and PSF1, a subunit of the GINS complex; and 2) proteins recruited to functional replication forks after origin firing during S phase, such as PCNA and Chk1 [23]. This analysis revealed that miR-28+ibrutinib combination treatment caused a specific reduction in the expression levels of proteins required for replication origin activation, including CDC45, CDC7, PSF1, PCNA, and Chk1, that were not observed in cells treated with miR-28 or ibrutinib individually (Figure 3D). Together, these results indicate that miR-28+ibrutinib combination therapy induces a specific cell cycle and DNA replication inhibitory transcriptional program in DLBCL cells.

### miR-28 potentiates the anti-tumor effect of ibrutinib against DLBCL by promoting cell cycle arrest

To analyze whether the transcriptional program induced by miR-28+ibrutinib combination therapy correlates with cell proliferation blockade, we first performed a pulsed BrdU and cell cycle analysis 20 hours after ibrutinib treatment in miR-28–transduced MD-901 ABC-DLBCL and SUDHL4 GCB-DLBCL cells. For each cell line, we used two ibrutinib doses (Figure 2A). When used individually as monotherapy, miR-28 and ibrutinib caused modest increases in the proportion of G_0_/G_1_ non-dividing cells and minor reductions in the proportion of BrdU-incorporating cells during S phase (Figure 4A). In contrast, cell cycle inhibition was greatly enhanced by miR28+ibrutinib combination (Figure 4A). Inhibition of DNA replication observed upon combined therapy was greater in MD-901 ABC-DLBCL cells than in SUDHL4 GCB-DLBCL cells (Figure 4A). The G2/M cell cycle phase was unaffected by either the individual therapies or the combined treatment. We also observed that miR-28 expression in ibrutinib-treated MD-901 cells increased the proportion of subG_0_/G_1_ dead cells detected 20 hours after ibrutinib administration (Figure 4B). Analysis of the Ki-67+ proliferative area of MD-901 ABC-DLBCL tumors in xenografts revealed that miR-28+ibrutinib combination therapy reduced proliferation (Figure 4C). Together, these results show that miR-28 potentiates the anti-tumor effect of ibrutinib by promoting DLBCL cell cycle arrest.

**Figure 4.**
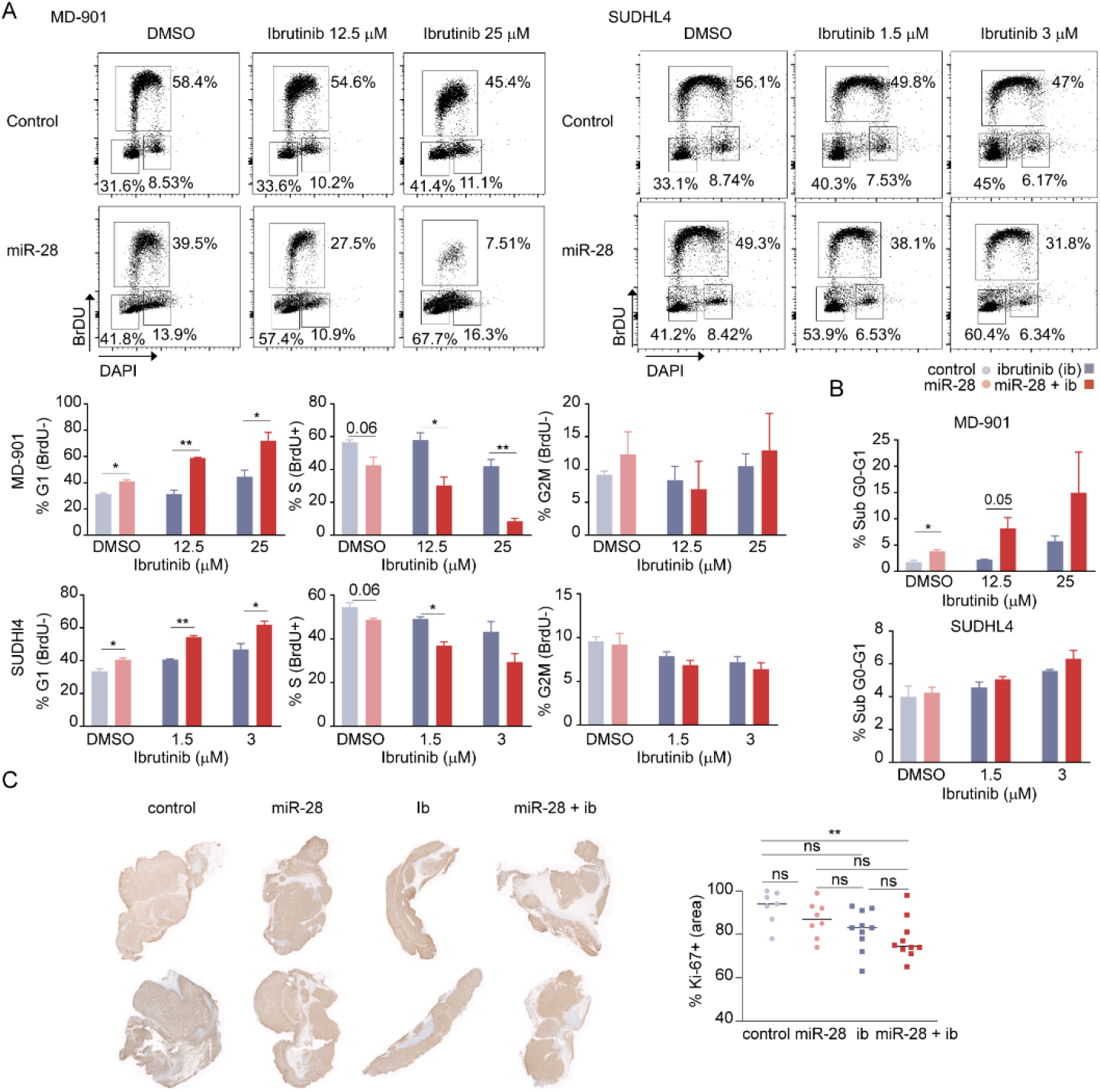
miR-28 potentiates the ibrutinib anti-tumor effect in DLBCL by promoting cell cycle arrest. **(A-B)** Cell cycle analysis in doxycycline-induced pTRIPZ-miR-28− or pTRIPZ-scramble–transduced MD-901 and SUDHL4 cells after 20 hours of ibrutinib treatment. **(A)** Representative flow cytometry plots of BrdU incorporation and cell cycle analysis, together with bar plots showing quantitative analysis of 2 independent experiments/cell line. *P<0.05, **P<0.01, unpaired *t* test. **(B)** Quantification of the DAPI-negative gated sub G_0_-G_1_ population. *P<0.05, unpaired *t* test. **(C)** Proliferation analysis of treated ABC-DLBCL tumors. Representative images are shown of Ki-67 immunohistochemistry in MD-901 tumors after miR-28 or ibrutinib monotherapy or miR-28+ibrutinib combination therapy. The graph shows quantification of Ki-67+ area **P<0.01, one-way ANOVA. Control (light gray), miR-28 (light pink), ibrutinib (dark gray), ibrutinib + miR-28 (dark red).

### miR-28+ibrutinib combination therapy impairs DNA replication

To investigate the molecular mechanisms underlying the S-phase inhibition induced by miR-28+ibrutinib combined treatment, we monitored DNA replication in a stretched DNA fiber assay after sequential pulse-labeling of ABC-DLBCL cells with thymidine analogs CIdU and IdU. This technique allows single-molecule resolution analysis of replicative dynamics through the quantification of fork progression rates and the frequency of active replication origins at a given time (Figure 5A; [24]). MD-901 cells transduced with inducible miR-28 or scramble were treated with or without ibrutinib for 20 hours. Both monotherapy treatments altered DNA replication dynamics (Figure 5B-D): miR-28 monotherapy caused a 50% reduction in the number of active origins and a concomitant two-fold increase in fork rate; in contrast, ibrutinib monotherapy decreased fork rate by 30% while increasing by 20% the number of active origins. Origin activity and fork rate are related parameters: alterations that slow down forks lead to the activation of more origins as a compensatory mechanism [25] [26], whereas alterations that primarily increase origin number lead to lower fork rates, likely due to competition for a limited dNTP pool [27] [28] [29]. However, this reciprocal association was disrupted in cells simultaneously exposed to miR-28 expression and ibrutinib. The reduction in the number of active origins associated with miR-28 treatment was maintained, but the compensatory increase in fork rate was reduced by 35% (Figure 5C, 5D). Therefore, miR-28+ibrutinib combination therapy limits DNA replication in DLBCL by simultaneously inhibiting origin activity and the compensatory acceleration of fork progression.

**Figure 5.**
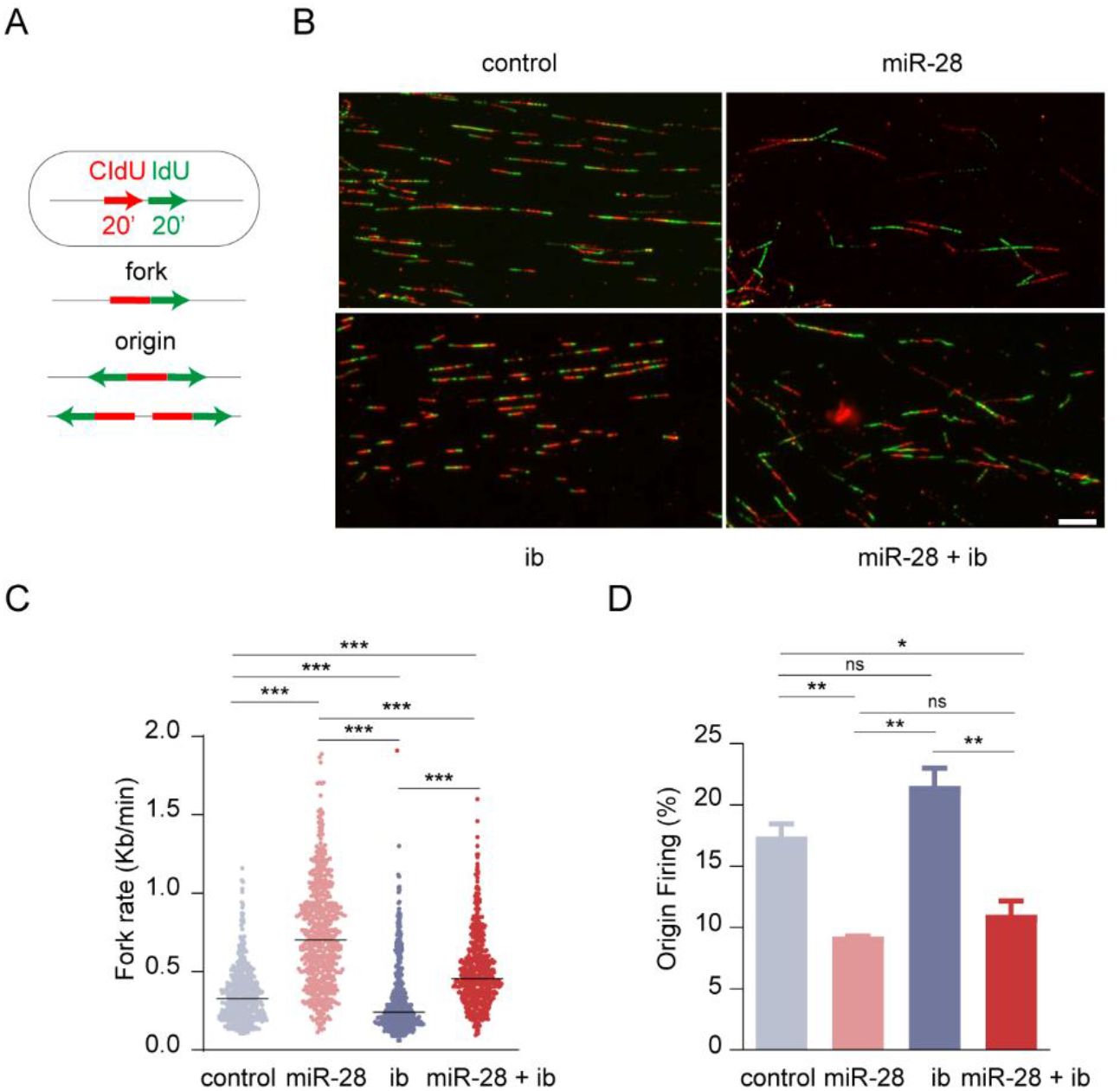
miR-28+ibrutinib combined treatment impairs DNA replication. **(A)** Experimental scheme of a stretched DNA fiber assay performed after 20 h exposure to 12.5 μM ibrutinib of doxycycline-induced pTRIPZ-miR-28 or pTRIPZ-scramble MD-901 cells. Cells were sequentially pulse-labeled with CIdU and IdU, and DNA replication patterns (fork rate and origin firing) were measured. **(B)** Representative images of DNA fibers after miR-28,ibrutinib monotherapy or combined treatment. **(C)** Dot plot showing the distribution of fork rate under the different conditions. Median values are depicted by horizontal black lines. The plots show data pooled from two replicates (n>100 cells per condition). ***P<0.001, one-way ANOVA. **(D)** Origin firing estimated as the percentage of first-label origin structures (n=2 independent experiments). *P<0.05, **P<0.01, one-way ANOVA.

### Survival of ABC-DLBCL patients is associated with reduced miR-28+ib signature

To assess whether the specific transcriptional program induced by miR-28+ibrutinib combination therapy is associated with survival differences in DLBCL patients, we ranked 62 ABC-DLBCL patients treated with R-CHOP immunochemotherapy and with 8 years of outcome data [4]. The ranking was based on a singscore analysis of the pretreatment expression of the 114 downregulated genes of the miR-28+ib signature [30] [31], such that a high score corresponds to higher expression and a low score to downregulated expression (Figure 6A). miR-28+ib signature scores were not associated with a particular ABC-DLBCL genetic subtype, and there were no significant differences in the proportions of BN2, MCD, and N1 genetic subtypes between patients with scores in the top (high expression) and bottom (low expression) tertiles (Figure 6B). Correlation with outcome data revealed that patients in the low-expression miR-28+ib signature tertile had higher overall survival than those in the high-expression tertile (p=0.005) (Figure 6C). These results indicate that the set of genes specifically downregulated by miR-28+ibrutinib combination therapy are an important determinant of ABC-DLBCL survival.

**Figure 6.**
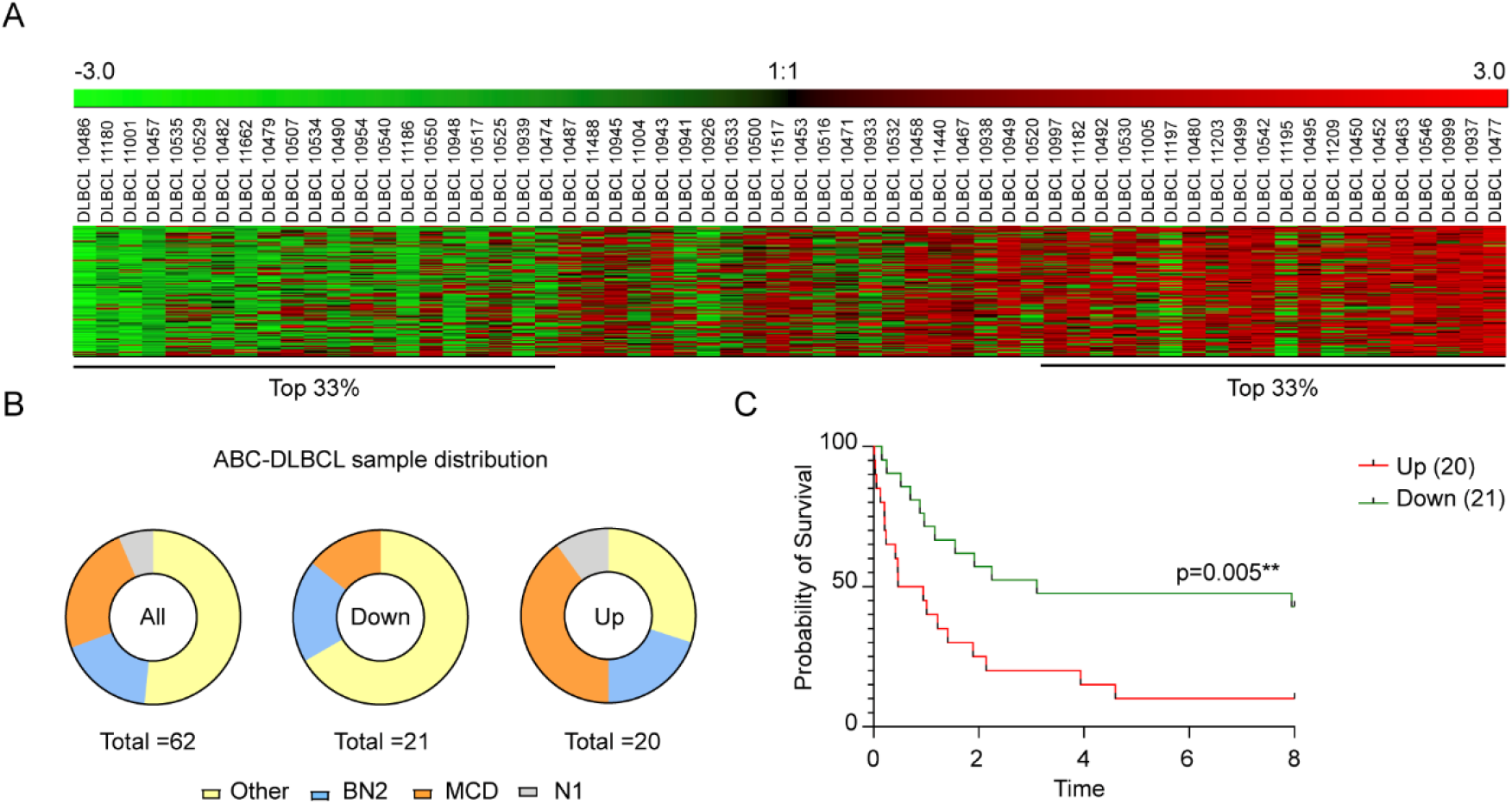
Downregulation of the miR-28+ib signature is associated with better survival in ABC-DLBCL patients. Survival analysis of ABC-DLBCL patients based on their scored expression of genes in the miR-28+ib signature. **(A)** Heatmap showing the scored gene expression profiles of 62 ABC-DLBCL patients before R-CHOP treatment for downregulated genes in the miR-28+ib signature, from the lowest expression score (left) to the highest (right). **(B)** Pie charts showing the distribution of ABC-DLBCL genetic subtypes (Other, BN2, MCD, N1) in all patients (left, n=62), in the 33% of patients with the lowest miR-28+ib signature expression (lowest tertile; middle, n=21), and in the 33% with the highest miR-28+ib signature expression (highest tertile; right, n=20). **(C)** Kaplan-Meier analysis of ABC-DLBCL patient survival stratified by scored miR-28+ib signature expression: highest expression tertile (Up) in red; lowest expression tertile (Down) in green. **P<0.01. Log-rank (Mantel-Cox) test.

## DISCUSSION

Despite the identification of molecular targets for DLBCL, a substantial proportion of these aggressive B cell lymphomas continues to present a major clinical challenge [9]. Here, we explored the therapeutic potential for DLBCL of the tumor suppressor miR-28, both as monotherapy and as combination therapy with ibrutinib. miR-28 expression is reduced in DLBCL and other forms of B-NHL [18, 19, 32] [33], and low miR-28 expression in DLBCL is associated with poor prognosis [34]. Using various ABC-DLBCL and GCB-DLBCL cell lines and PDX phase-II preclinical xenograft models [21], we found that miR-28 inhibits DLBCL growth *in vivo*. Transcriptome profiling of miR-28–treated PDX DLBCL tumors showed downregulation of genes involved in tumorigenesis. In an earlier study, we reported that miR-28 downregulates B-cell expression of a gene network downstream of the BCR signaling pathway and that miR-28 expression correlates inversely with that of genes downstream of BCR signaling, including NF-κB2, IKKB, and Bcl-2 in ABC-DLBCL[19]. Several of the genes targeted by miR-28 in B cells, including Bcl-2, NF-κB2 and IRAK1, have been shown to be essential for oncogenic signaling in the ABC and GCB genetic subtypes of DLBCL [35], and our findings are thus in agreement with the anti-tumor effect of miR-28 observed in ABC-and GCB-DLBCL.

Combination therapies are a foundation of cancer treatments because the simultaneous use of two or more agents can target key pathways in a synergistic or additive manner [36]. Ibrutinib monotherapy is initially effective in a high fraction of patients with ABC-DLBCL, but resistance to BTK inhibitors develops rapidly, even in tumors addicted to BCR-dependent NF-*k*B activation, and progression-free survival is therefore very short (2 months) [22, 37]. Ibrutinib combination regimes are currently under exploration [11], and ibrutinib plus R-CHOP chemotherapy has yielded excellent results, but only in young ABC-DLBCL patients with the MCD and N1 genetic subtypes [13]. Therefore, finding new ibrutinib combination therapies is a crucial clinical need for the treatment of DLBCL.

miRNA-based therapies are especially attractive for cancer and B cell neoplasia because of their potential to simultaneously inhibit multiple targets, which limits the possibility of treatment failure due to tumor evolution through alternative oncogenic pathways and the selection of clones with a treatment-resistant mutation [16, 17]. To our knowledge, this study is the first to assess a miRNA-based combination therapy with ibrutinib. Using *in vitro* assays and *in vivo* xenograft models combined with RNA-Seq transcriptome profiling, pulsed BrdU incorporation, and DNA fibers assays, we found that miR-28 enhances the anti-tumor effect of ibrutinib by inhibiting a DNA replication and cell cycle transcriptional gene network in DLBCL cells.

The inhibition of this proliferation network is specific to miR-28+ibrutinib combination therapy and is not induced by monotherapy with either agent. This suggests that the molecular actions of miR-28 and ibrutinib intertwine synergistically to impair DNA replication and cell division in DLBCL cells. Of note, poor DLBCL outcomes have been linked to high expression of proliferation genes and a high Ki-67 proliferation index [38] [39] [40]. Likewise, high expression of proliferation genes is considered one of the main oncogenic signatures of DLBCL[4]. Thus, our finding that the unique set of genes downregulated by miR-28+ibrutinib combination therapy is also downregulated in ABC-DLBCL patients with better survival, strongly supports that this molecular pathway is key to disease outcome. We propose that the miR-28+ib signature described here can potentially be a valuable new gene expression panel for ABC-DLBCL prognosis and stratification.

Regarding the therapeutic implications, our study provides evidence for the effectiveness of a new miRNA-based ibrutinib combination therapy for DLBCL. The correlation analysis of miR-28+ib signature expression with clinical outcome in an R-CHOP-treated ABC-DLBCL cohort suggests that miR-28+ibrutinib combination therapy would benefit patients with any ABC-DLBCL genetic subtype. Future studies focused on: i) B-cell lymphoma specific miR-28+ibrutinib delivery approaches and ii) analyzing if miR-28+ibrutinib combination therapy reduces the generation of ibrutinib-resistant DLBCL clones, will further increase the translational potential of this work. Interestingly, we have found that miR-28 targets 61 genes essential for ibrutinib-resistant ABC-DLBCL growth [41], including Bcl-2, ORC6, and IRAK1, suggesting that miR-28+ibrutinib combination therapy could reduce the generation of ibrutinib resistance and provide a more durable treatment for DLBCL patients.

In summary, the results of this study provide insights for the development of new diagnostic and therapeutic strategies based on miR-28 plus ibrutinib combination.

## MATERIALS AND METHODS

### Cell culture

Human DLBCL cell lines (DOHH2, DB, OCI Ly19, MD-901, RIVA, U2932, HBL1, and SUDHL4) were grown at 37°C in the presence of 5% CO_2_ and maintained in RPMI supplemented with 10% fetal bovine serum, 1% penicillin/streptomycin, and 10 mM Hepes (Gibco). All cell lines were regularly tested for mycoplasma with the MycoAlert PLUS Mycoplasma Detection Kit (Lonza).

### Mice

Nonobese diabetic severe combined immunodeficiency G (NOD/SCID/IL-2rγnull; NSG) mice were housed in the Centro Nacional de Investigaciones Cardiovasculares (CNIC) animal facility under specific pathogen-free conditions. All animal procedures conformed to EU Directive 2010/63EU and Recommendation 2007/526/EC regarding the protection of animals used for experimental and other scientific purposes, enacted in Spanish law under RD 53/2013. Animal procedures were reviewed by the CNIC Institutional Animal Care and Use Committee (IACUC) and approved by the Consejeria de Medio Ambiente, Administración Local y Ordenación del Territorio of the Comunidad de Madrid.

### Lymphoma models

For subcutaneous (SC) xenografts, 1–5 × 10^6^ DOHH2, DB, OCI Ly19, MD-901, or RIVA cells were mixed in a 1:1 in Matrigel (BD Biosciences)/PBS and injected into the flanks of 10-week-old NSG mice. In mice injected with transduced MD-901 cells, doxycycline (Sigma-Aldrich) was administered twice per week at 0.04% in the drinking water, and mice received daily 40 mg/kg intraperitoneal (IP) injections of ibrutinib (PCI-32765) (MedChemExpress) from day 10 after the SC xenograft. Ibrutinib was dissolved in 10% DMSO and 90% corn oil, and 200 ul of this solution was administered to each mouse. Untreated groups received IP injections of the 10% DMSO, 90% corn oil solution.

To establish subcutaneous DFBL-18689-V2 PDX tumors, NSG mice received intravenous (IV) injections of DFBL-18689-V2 PDX cells, and splenocytes were extracted after 23-28 days. 1-3×10^6^ cells were resuspended in PBS, mixed with 30%-50% matrigel, and injected SC into the flanks of 6-8-week-old NSG.

Subcutaneous tumor volume was measured with a digital caliber using the following formula: volume = (width)^2^ x length/2. Tumor growth was calculated for each individual tumor as the tumor volume at the indicated time point normalized to the initial tumor volume (at the time of starting treatment).

### Treatment with miR-28 mimics

miRNA mimics of miR-28 and the scramble control sequence were purchased from Ambion. Established DFBL-18689-V2 PDX tumors (> 100 mm^3^) or DLBCL tumors (> 150 mm^3^) were treated with weekly intratumor injections of 0.5 nmol miRNA mimics (miR-28 or control) together with invivofectamine (Ambion).

### FACS analysis of DFBL-18689-V2 PDX cells

DFBL-18689-V2 was acquired from the Public Repository of Xenografts (PRoXe). PDX cells (0.25–1×10^6^) were injected IV into 6-8-week-old NSG mice, and mice were sacrificed on day 26-31 after injection. Single-cell suspensions were obtained from spleens, and erythrocytes were lysed (ACK Lysing Buffer). Cells were stained with combinations of the following antibodies: anti-mouse CD45-APC, anti-mouse CD45-V450, anti-human IgM-APC, anti-human CD20-FITC, and anti-human CD19-APC (all from BD Pharmigen). For proliferation analysis, cells were fixed and permeabilized using the Intracellular Fixation and Permeabilization Buffer Set (eBiosience) and stained with anti-ki67 (Abcam); staining was revealed with alexa fluor 647 goat anti-rabbit igG (H+L) (L.T.). Live cells were detected with DAPI (Sigma-Aldrich) or with LIVE/DEAD Fixable Yellow Dead Cell Stain (Thermo Fisher). Labeled cells were acquired with a LSRFortessa High-Parameter Flow Cytometer and analyzed with FlowJo V10.4.2 software.

### DFBL-18689-V2 PDX transcriptome analysis

DFBL-18689-V2 PDX SC tumors were treated by intratumoral injection with 0.5 nmol of miR-28 or control mimics. Three independent pairs of DFBL-18689-V2 PDX tumors treated with miR-28 or control obtained at day 5 after the mimic treatment were used for RNA-seq sequencing. Total RNA was used to generate barcoded RNA-seq libraries using the NEBNext Ultra RNA Library preparation kit (New England Biolabs). First, poly A+ RNA was purified using poly-T oligo-attached magnetic beads followed by fragmentation and first and second cDNA strand synthesis. Next, cDNA ends were repaired and adenylated. The NEBNext adaptor was then ligated followed by Uracile excision from the adaptor and PCR amplification. Libraries were sequenced on a HiSeq2500 (Illumina) to generate 60 bases single reads. FastQ files for each sample were obtained using CASAVA v1.8 software (Illumina). Differential expression between miR-28- and control-treated tumors was analyzed by ANOVA (314 protein coding transcripts, p-value <0.05), and expression values were presented in a heatmap.

Alterations to disease and functions in miR-28- and control-treated tumors were identified by Ingenuity pathway analysis (IPA) of differentially expressed genes (DEGs) (p-value < 0.05).

### Gene set enrichment analysis and gene signature analysis

The miR-28–induced transcriptome profile in the DFBL-18689-V2 PDX model was compared with the DLBCL gene signature through a Geo2R analysis of raw data from GEO GSE44337 [42]. For the differential expression analysis between DLBCL and PB B-cells, we used DLBCL samples (GSM1083475, GSM1083476, GSM1083477, GSM1083478, GSM1083479, GSM1083480, GSM1083481, GSM1083482, and GSM1083483) and human peripheral blood B-cells (GSM1083472, GSM1083473, and GSM1083474). Data were filtered for an adjusted p-value < 0.05 and logFC < −1.5 or >1.5, and the DLBCL gene signature was obtained as logFC >1.5. Enrichment of the DLBCL gene signature in the RNA-Seq data from miR-28–treated PDX tumors was tested by Gene Set Enrichment Analysis (GSEA).

For a more specific characterization of the pathways altered by the combined action of miR-28 and ibrutinib in MD-901 cells, GSEA was performed to determine whether there was enrichment of Reactome gene sets. The rank list used for this analysis was the DEG analysis comparing miR-28+ibrutinib versus control.

ABC-DLBCL patients with outcome data [4] were ranked using singscore [30] [31] according to their enrichment score for the miR-28+ib signature (downregulated genes) before R-CHOP treatment.

### Lentiviral expression constructs

Lentiviral constructs were generated to transduce the human DLBCL cell lines HBL1, MD-901, U2932, and SUDHL4. The miR-28 precursor sequence was cloned into the doxycycline-inducible pTRIPZ vector (Thermo scientific), and a scramble pTRIPZ vector was used as a control. Lentiviral supernatants were obtained 48 hours after transfection of 293T cells with packaging vectors (VSVG, Δ9.8) and pTRIPZ plasmids. Transduced human lymphoma cell lines were selected by culture in the presence of 0.4 μg/ml puromycin for 3 days. Expression of miR-28 or the scramble control sequence was induced by exposure to 0.5 μg/ml doxycycline for 2 days. Transduced RFP+ cells were isolated with a FACS Aria cell sorter for subsequent experiments.

### miR-28+ ibrutinib combined treatment analysis *in vitro*

Human DLBCL cells (HBL1, MD-901, U2932 and SUDHL4) transduced with pTRIPZ-miR-28 or pTRIPZ-scramble were counted, and 200,000 cells were seeded in duplicate on a 48-well plate in fresh medium containing doxycycline at 0.5 μg/ml to induce miRNA expression. Serial dilutions of ibrutinib (Selleck Chem) were prepared in vehicle (DMSO), and equal volumes of the diluted drug were added to cells to achieve the desired final concentration. At least two independent experiments were performed per cell line. The number of viable cells was determined 72 hours after the start of ibrutinib treatment using CellTiter-Glo® reagent. Luminescence was measured with an Orion Plate Luminometer reader (Berthold). Luminescence values in medium-only wells were subtracted to yield relative luminescence units (RLU), and the data were normalized to DMSO-treated or ibrutinib-treated control cells. The percentage of viable cells was measured 72 hours after ibrutinib treatment by DAPI staining (with viable cells gated as DAPI-), and RFP expression was measured by flow cytometry in an LSRFortessa High-Parameter Flow Cytometer and analyzed with FlowJo V10.4.2 software.

### MD-901 miR-28+ibrutinib combined treatment transcriptome analysis

MD-901 cells transduced with pTRIPZ-scramble or pTRIPZ-miR-28 were induced with doxycycline (0.5 μg/ml) for 24 hours. Cells were then seeded at 200,000 cells/ml and treated with 25 μM ibrutinib, while maintaining doxycycline in the medium. As a control, cells were treated with vehicle (DMSO). RFP+ cells were isolated in a FACS Aria cell sorter at 0, 4, and 20 h after the start of ibrutinib treatment. RNA-Seq analysis was performed on two independent combined-treatment experiments. Total RNA was used to generate barcoded RNA-seq libraries using the NEBNext Ultra II Directional RNA Library preparation kit (New England Biolabs) according to manufacturer’s instructions. First, poly A+ RNA was purified using poly-T oligo-attached magnetic beads followed by fragmentation and first and second cDNA strand synthesis. Next, cDNA ends were repaired and adenylated. The NEBNext adaptor was then ligated followed by second strand removal, uracile excision from the adaptor and PCR amplification. Libraries were sequenced on a HiSeq4000 (Illumina) to generate 60 bases single reads. FastQ files for each sample were obtained using bcl2fastq 2.20 Software (Illumina). Differential expression analysis was performed with the Limma R package at the different time points (0, 4, and 20 hours) and also time-independently.

### miR-28+ib signature

The miR-28+ib signature was obtained through comparison of RNA-Seq data from pTRIPZ-transduced MD-901 cells expressing miR-28 or the control scramble sequence and exposed to vehicle (DMSO) or ibrutinib for 20 hours, yielding three experimental conditions: single treatment with miR-28, single treatment with ibrutinib, and combined treatment with miR-28+ibrutinib. All analyses were filtered for an adjusted p-value < 0.05. Three overlap analyses were performed to obtain the miR-28+ib signature, defined as genes that changed with the combined treatment but not with the single treatments. First, to exclude genes that change with ibrutinib treatment, an overlap analysis was performed between DEGs obtained with the comparison of miR-28+ibrutinib versus miR-28 and ibrutinib versus control; since there was no overlap, the whole set of 455 genes was included in ‘list 1’. Second, to exclude genes that change with miR-28 treatment, an overlap analysis was performed between DEGs obtained with the comparison of miR-28+ibrutinib versus ibrutinib and miR-28 versus control; the overlapping 3290 genes were eliminated, and the remaining 1437 genes were assigned to ‘list 2’. Third, an overlap analysis was performed between list 1 and list 2, and the resulting list of 157 overlapping genes was defined as the miR-28+ib signature (Figure S3B).

GO terms were assigned to each gene in the signature. A heatmap was generated to represent the Z-score expression values of the 157 genes in miR-28+ib signature for the different treatments and time points, and the 7 most representative GO terms were depicted.

Ingenuity Pathway Analysis (IPA) was used to identify the distinct signaling pathways and networks in miR-28+ib signature that differed between the miR-28+ibrutinib and control (untreated) conditions. Two merged DNA replication-related networks identified with IPA were represented using STRING tool.

### Comparative Ingenuity Pathway Analysis

To analyze signaling pathways altered by the single and combined treatments, a comparative IPA was performed with the following three time-independent differential expression analysis (filtered for an adjusted p-value <0.05 and logFC <-1 or > 1): miR-28 versus control, ibrutinib versus control, and miR-28+ibrutinib versus control.

### Cell proliferation

MD-901 and SUDHL4 cells transduced with pTRIPZ-scramble or pTRIPZ-miR-28 were induced with doxycycline (0.5 μg/ml) for 24 hours. Cells were then plated at 200,000 cells/ml and treated with the indicated ibrutinib concentrations in medium containing doxycycline. After 20 hours, the cell cycle was analyzed using DAPI staining and the FITC BrdU Flow kit (BD Pharmigen). The sub G0-G1 population was determined in DAPI-negative gated cells.

### Histopathology

Tumors were fixed in formalin and embedded in paraffin. Sections were cut and stained with hematoxylin and eosin (Sigma) or anti-Ki-67 (rabbit polyclonal antibody, Life Technologies, Thermofisher, PA5-19462). Ki-67 positive lymphoma areas were quantified on scanned slides using NDP.view2 plus software.

### Protein detection by immunoblot

Control and miR-28-pTRIPZ-transduced MD-901 cells were induced with doxycycline, plated at 200,000 cells/ml, and treated for 20 h with 25 μM ibrutinib with doxycycline in the medium. Whole cell extracts were prepared by cell resuspension in Laemmli buffer (50 mM Tris–HCl pH 6.8, 10% glycerol, 3% SDS, 0.006 w/v bromophenol blue, 5% 2-mercaptoethanol) followed by sonication (3 × 4 sec) in a Branson Digital Sonifier set at 50% amplitude. Immunoblots were performed using standard protocols. Primary antibodies used in this study: anti-CDC45 (generated in Méndez Lab, 1:1,000; [43]), anti-CDC7 (Novus Biologicals NB120-17880, 1:1,000), anti-PCNA (Proteintech 60097-1-Ig, 1:1,000), anti-PSF1 (generated in Méndez Lab, 1:1,000; [43]), anti-CHK1 (Santa Cruz Biotechnology sc-8408; 1:10,000), anti-H3 (Abcam ab1791, 1:20,000). LI-COR secondary antibodies: IRDye 800CW goat anti-rabbit (926-32211) and IRDye 680RD goat anti-mouse (926-68070; LI-COR, Bad Homburg, Germany).

### Single-molecule analysis of DNA replication

Control and miR-28 pTRIPZ-transduced MD-901 cells were induced with doxycycline, plated at 300,000 cells/ml, and treated for 20 h with 12.5 μM ibrutinib and doxycycline in the medium. Cells were pulse-labeled sequentially with 50 μM 5-chloro-2’-deoxyuridine (CldU; 20 min) and 250 μM 5-iodo-2’-deoxyuridine (IdU; 20 min). Labeled cells were lysed in 0.2 M Tris pH 7.4, 50 mM EDTA, 0.5% SDS. Stretched DNA fibers were prepared as described (Terret et al, 2009). For the immunodetection of labeled tracks, fibers were incubated with primary antibodies anti-CldU (rat monoclonal anti-BrdU, Abcam, AB6326), anti-IdU (mouse monoclonal anti-BrdU, BD Biosciences, 347580) and anti-ssDNA (mouse monoclonal IgG2a, Millipore MAB3034) for 1 h at RT in a humidity chamber, followed by incubation with Alexa Fluor-conjugated secondary antibodies (Invitrogen/Molecular Probes, A-11007, A-21121, and A-21241) for 30 min at RT. Microscopy images were obtained with a DM6000 B Leica microscope equipped with an HCX PL APO 40×, 0.75 NA objective. A standard conversion rate of 1 μm=2.59 kb was used [44]. Signals were measured and quantified using ImageJ software. For the analysis of fork rate, the length of the green track was measured in red–green structures (>300 tracks measured per condition and replicate). The frequency of origin activation was determined as the percentage of green–red–green structures (origins activated during the CldU pulse) relative to all replicative structures containing a red track, as described [24]. >500 structures were measured per condition and replicate. A conversion factor 1 μm = 2.59 kb was used [44].

### Statistics

Statistical analyses were performed in GraphPad Prism 8. Error bars represent standard deviation of the mean. Data were analyzed using two-tailed unpaired Student t-*test* for comparison of two experimental groups or ANOVA for comparison of more than two groups. For xenograft experiments comparing two or more groups over time, linear mixed models were run with R software. Differences were considered significant at p-values < 0.05.

## ACKOWLEDGEMENTS

We thank Simon Bartlett for English editing. T.F.N. and P.U.-C. were supported by PhD fellowships from the Spanish Ministerio de Ciencia, Innovacion y Universidades (BES-2017-079759 and BES-2017-081887, respectively), CG-E by a PhD fellowship awarded by La Caixa España 2017, AS-N by a FPI Severo Ochoa fellow (PRE2018-083475), E.A.C. by funding from the Comunidad de Madrid (CT4/21/PEJ-2020-AI/BMD-18112), A.R.R. by CNIC funding and V.G.Y. by the Universidad Complutense de Madrid. This work was funded by Spanish Ministerio de Ciencia e Innovación grants PID2019-107551RB-I00/AEI/10.13039/501100011033 to V.G.Y., PID2019-106773RB-I00/ AEI / 10.13039/501100011033 to A.R.R. and PID2019-106707RB-100/AEI/10.13039/50110001103v to J.M, co-sponsored by EU ERDF funds. The CNIC is supported by the Instituto de Salud Carlos III (ISCIII), the Ministerio de Ciencia e Innovación (MCIN) and the Pro CNIC Foundation), and is a Severo Ochoa Center of Excellence (grant CEX2020-001041-S funded by MICIN/AEI/10.13039/501100011033).

## AUTHOR CONTRIBUTIONS

T.F., E.A.C., and P.U-C performed experiments. A.S.N. and C.G.E. performed computational analysis. T.F., E.A.C, P.U, A.M. and V.G.Y. analyzed and interpreted data. T.F prepared figures. J.M. contributed to the conception of the experimental design. V.G.Y and A.R.R. conceived the overall strategy. V.G.Y, T.F and A.R.R wrote the manuscript. All authors revised and contributed to the final manuscript.

