## Supplementary Figures for "miR-28 plus ibrutinib as a novel combination therapy for Diffuse Large B Cell Lymphoma"

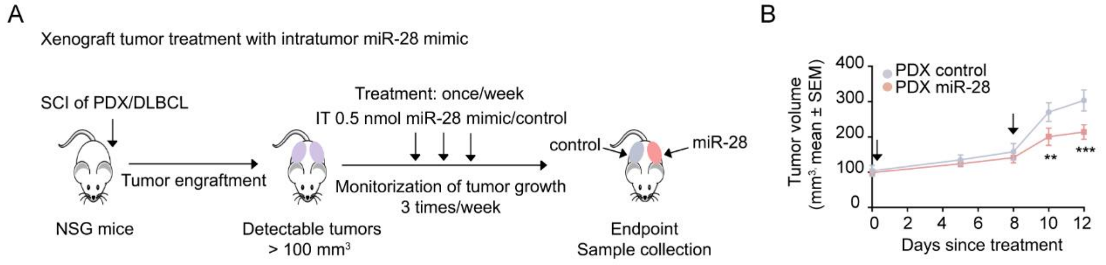


Figure S1

**Figure S1. DLBCL xenografts models treated by intratumoral injection of miR-28 mimics. (A)** Experimental model. DLBCL cell lines or ABC-DLBCL PDX (PRoXe DFBL-18689-V2) cells were injected subcutaneously into NSG mice. When tumors were detectable (>150 $mm^3$ for DLBCL cell lines and > 100 $mm^3$ for ABC-DLBCL PDX), mice were given intratumoral injections of 0.5 nmol miRNA mimics: miR-28 (pink) or control (gray). Tumor volume was measured 3 times per week. **(B)** Volume of ABC-DLBCL PDX tumors at the end of treatment, calculated as mean volume ± SEM (3 independent experiments, n=22 control-treated tumors, n=21 miR-28-treated tumors). **P<0.01, ***P<0.001, linear mixed model.

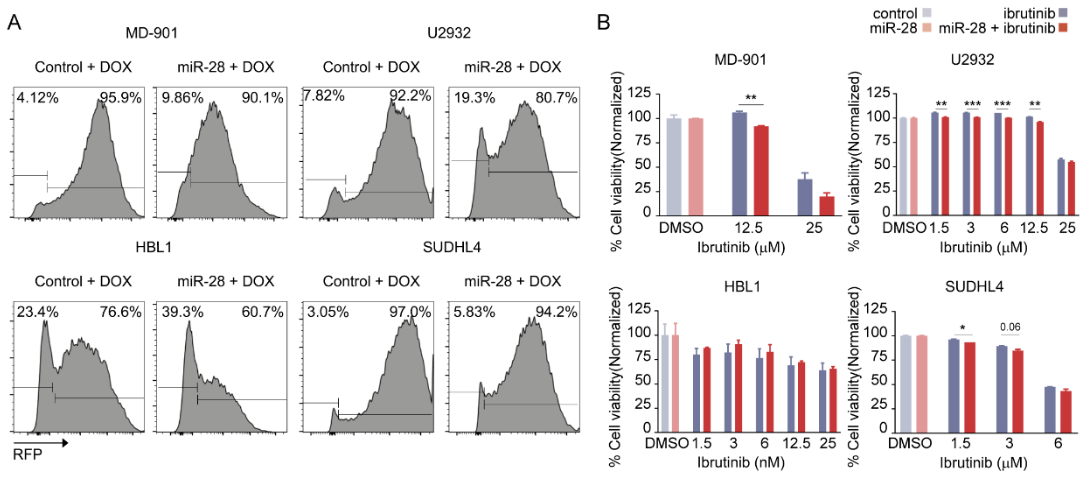


Figure S2

**Figure S2. RFP expression and viability of DLBCL cell lines after miR-28+ibrutinib combined treatment. (A)** Representative flow cytometry plots of RFP expression by pTRIPZ-miR-28– or pTRIPZ-scramble–transduced DLBCL cell lines induced with doxycycline for 3 days. **(B)** pTRIPZ-miR-28– or pTRIPZ-scramble–transduced DLBCL cell lines were treated with serial dilutions of ibrutinib in the presence of doxycycline. The percentage of viable cells was analyzed at day 3 by FACS with DAPI staining (viable cells gated as DAPI negative). The bar plots show quantification of cell viability in response to miR-28+ibrutinib combined treatment (dark red) normalized to miR-28 treatment (light pink) and in response to ibrutinib treatment (dark gray) normalized to control (light gray). *P<0.05, **P<0.01, ***P<0.001, unpaired *t* test.

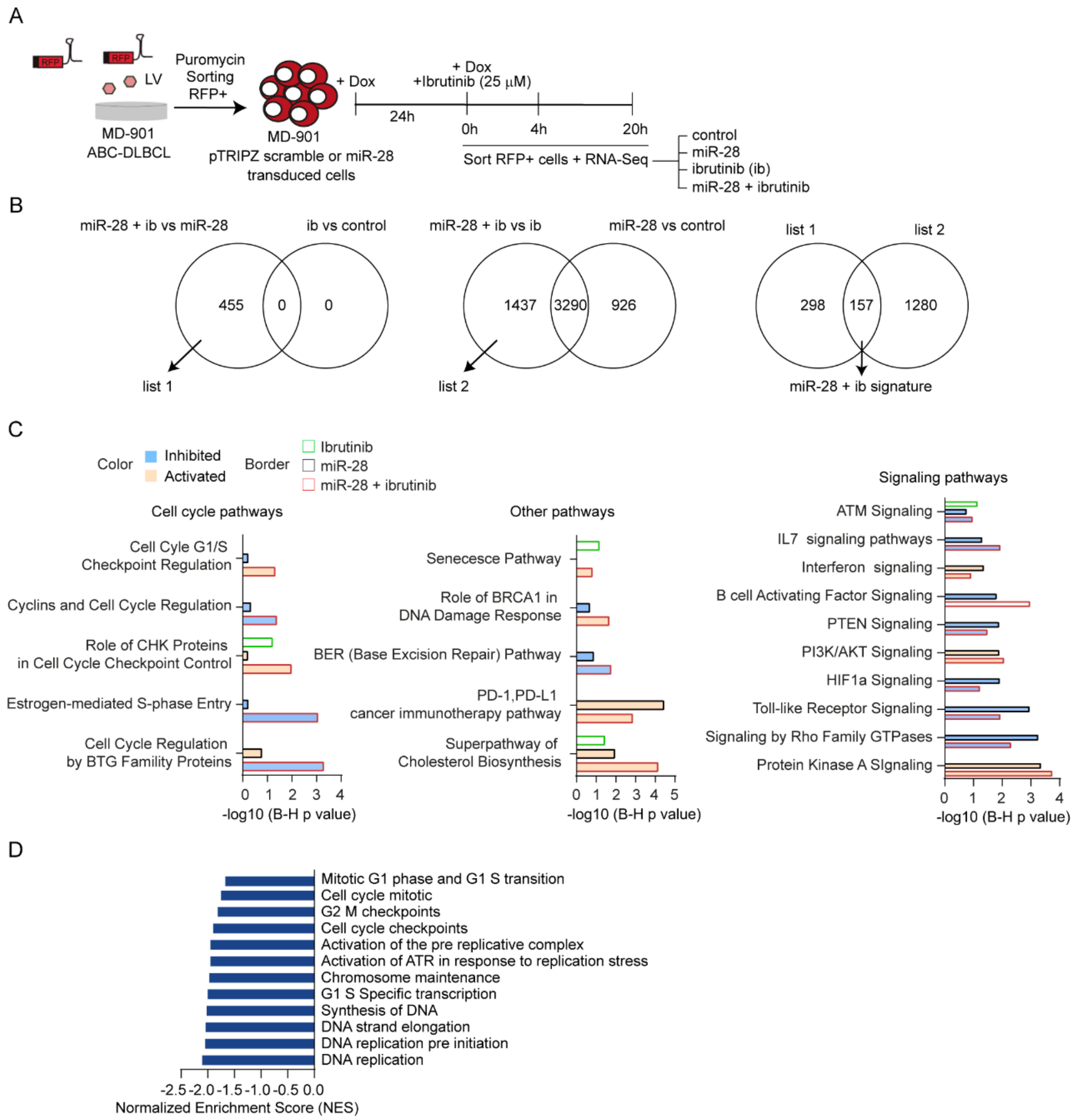


**Figure S3**

**Figure S3. Effect of miR-28+ibrutinib combined treatment on the ABC-DLBCL transcriptome. (A)** Experimental setup. pTRIPZ-miR-28– or pTRIPZ-scramble–transduced MD-901 cells were selected in the presence of puromycin, and RFP$^+$ cells were isolated by FACS. Cells were induced with doxycycline for 24 hours before treatment with ibrutinib (25 µM). RNA-Seq was performed in RFP$^+$ cells isolated by FACS after 0, 4, and 20h ibrutinib treatment. **(B)** Analysis used to identify the miR-28+ib signature. **(C)** Bar plots showing Benjamini-Hochberg (B-H) adjusted p-values (q val) for comparative Ingenuity Pathway Enrichment Analysis of differentially expressed genes

(DEGs) in response to ibrutinib (green border), miR-28 (black border), and miR-28+ibrutinib (red border) compared with control. Terms are colored according to the z-score (pathway activation prediction): positive values (activation in orange) and negative values (inhibition in blue). **(D)** Bar graph showing Normalized Enrichment Scores (NES) for the gene set enrichment analysis of miR-28+ibrutinib combined treatment versus control.
